# Human auditory cortex preferentially tracks speech over music without explicit attention

**DOI:** 10.64898/2026.03.12.710296

**Authors:** Rajvi Agravat, Maansi Desai, Alyssa M. Field, Sandra Georges, Jacob Leisawitz, Gabrielle Foox, Saman Asghar, Dave Clarke, Elizabeth C. Tyler-Kabara, M. Omar Iqbal, Andrew J. Watrous, Anne E. Anderson, Howard L. Weiner, Liberty S. Hamilton

## Abstract

Our brains constantly filter incoming sounds to understand our environment. While extensively studied in adults, how this ability develops across childhood remains unclear. We recorded intracranial brain activity from 54 participants aged 4-21 while they watched movie clips containing simultaneous speech and music. We used deep neural networks to separate the mixed audio into isolated speech and music streams, then built encoding models to determine which stream best predicted neural responses in the auditory cortex. Although participants heard only the original mixture with no instruction to attend to either stream, higher-order auditory regions including the superior temporal gyrus (STG), superior temporal sulcus (STS) and middle temporal gyrus (MTG), responded preferentially to speech. This speech-bias strengthened with age in STG, suggesting that this region progressively sharpens its representation of socially relevant sound across development. These findings indicate that speech prioritization in the developing brain emerges automatically, without directed attention.

## Introduction

The human auditory system is adept at disentangling environments where multiple sound sources, like speech and music, coexist and overlap. This process, known as auditory streaming, enables listeners to perceive relevant sounds amid noise (Cherry, 1953; Shinn-Cunningham, 2008; Zatorre, Bouffard, et al., 2002), allowing us to follow conversations in noisy settings or appreciate music in a crowded venue. Yet, despite extensive research on auditory processing in adults, two key questions remain unresolved. First, how does the auditory cortex process overlapping speech and music when they co-occur in naturalistic contexts? Do certain regions respond preferentially to one stimulus type over the other, without explicit attention? Second, how do the brain circuits supporting auditory streaming develop across childhood, adolescence, and adulthood?

Recent computational and systems neuroscience research has deepened our understanding of how subregions of the auditory cortex respond specifically to speech versus music but mostly using isolated and nonoverlapping sounds. When tested with isolated sounds, some groups have shown that superior and middle temporal gyri (STG and MTG) exhibit stronger responses to speech, while regions such as planum polare (PP) and anterior STG show stronger responses to music (Angulo-Perkins et al., 2014; Belin et al., 2000; Boebinger et al., 2021; McCarty et al., 2023; S. Norman-Haignere et al., 2015; S. V. Norman-Haignere et al., 2022). However, other evidence suggests more distributed representations across cortical regions. For example, with domain-selective responses restricted to frequency-specific oscillatory patterns rather than anatomically distinct areas (te Rietmolen et al., 2024). Alternatively, selectivity for music has been observed in both anterior and posterior STG depending on the specific acoustic features and task demands (S. V. Norman-Haignere et al., 2022). These patterns reflect a broader functional specialization within the auditory cortex. Electrophysiological studies further reveal that the posterior and middle STG exhibit robust tuning to phonetic and syntactic features of speech across both single-talker (Bhaya-Grossman & Chang, 2022; de Heer et al., 2017; Hamilton et al., 2018, 2021; Howard et al., 2000; Hullett et al., 2016; Mesgarani et al., 2014) and multi-talker contexts (Mesgarani & Chang, 2012; Raghavan et al., 2023). In contrast, music cognition studies show preferential responses to musical pitch, harmony, and melody in bilateral STG, posterior MTG, and auditory association cortex (Angulo-Perkins et al., 2014; Hugdahl et al., 1999; McCarty et al., 2023; Peretz, 2006). Recent intracranial studies examining cortical tracking during naturalistic listening to alternating speech and music reveal that speech is represented in the perisylvian language regions, and music in superior temporal, middle temporal, and supramarginal regions (Osorio & Assaneo, 2025), representing an important step toward understanding speech-music distinctions in more natural contexts.

Despite this progress, most research characterizing speech-music distinctions has relied on isolated, non-overlapping stimuli, leaving open fundamental questions about how the auditory cortex manages competing incoming streams. Speech and music share fundamental acoustic features including pitch, rhythm, and hierarchical structure (Koelsch et al., 2013; A. D. Patel, 2003; Sankaran et al., 2024), and engage overlapping cortical areas. Yet subtle differences in how these features are weighted can lead to distinct activation patterns within shared networks (Abrams et al., 2011; S. Norman-Haignere et al., 2015). Furthermore, while these distinctions are well characterized in adults, much less is known about how such functional organizations emerge and evolve over the course of development. Behavioral and electrophysiological studies indicate that auditory stream segregation abilities emerge early but mature progressively throughout childhood and adolescence (Jones et al., 2015; Leibold, 2019; Vander Ghinst et al., 2019), yet direct neural evidence of speech versus music selectivity during development, under naturalistic listening conditions, remains limited.

To fill this gap, we collected intracranial stereo-electroencephalography (sEEG) recordings from children, adolescent, and young adult epilepsy patients listening to naturalistic audiovisual stimuli (movie clips). sEEG offers a unique opportunity to study the neural representation of overlapping speech and music with high spatial and temporal resolution (Berezutskaya et al., 2017; Chang, 2015; Flinker et al., 2011; Hamilton et al., 2021; Howard et al., 2000; Lachaux et al., 2012; Mercier et al., 2022; Micheli et al., 2020; Ozker et al., 2017), and provides basic science insights into local circuit dynamics that are inaccessible to noninvasive methods (Chang 2015; Mercier et al. 2022). Using encoding models to predict neurophysiological data, we investigate whether such selectivity emerges during passive listening to mixed auditory scenes in the developing brain, and how this selectivity evolves from childhood to young adulthood. This research advances our understanding of competing sound perception in the developing brain, in the absence of explicit attention.

## Results

Understanding how the brain encodes meaningful sounds in the presence of competing auditory information is critical for explaining one of the most impressive feats of human perception: following a behaviorally relevant auditory stream amid background sounds. The goal of our study was to uncover how the developing auditory system handles this challenge. Specifically, we asked how acoustic features of speech and music are encoded in the brain even when heard simultaneously and without explicit attention directed to either stream.

To address this question, we recorded sEEG from 54 pediatric, adolescent, and young adult epilepsy patients (29M/25F; ages 4-21; mean 12.59 ± 4.7 years) undergoing surgical monitoring To address this question, we recorded sEEG from 54 pediatric, adolescent, and young adult to treat drug resistant epilepsy (Figure 1A). Participant ages spanned four development stages: early childhood (ages 4-5, n=4), middle childhood (ages 6-11, n=22), early adolescence (ages 12-17, n=17), and late adolescence (ages 18-21, n=11) (Figure 1B). Patients listened to movie trailer clips containing overlapping speech and music (Figure 1C) while we recorded brain activity across the left and right auditory cortex. This included 9,874 electrode contacts across all patients, out of which 1,161 contacts were distributed across the temporal lobe areas including Heschl’s gyrus (HG), planum temporale (PT), and planum polare (PP), superior temporal gyrus (STG), superior temporal sulcus (STS), and middle temporal gyrus (MTG) bilaterally (Figure 1A-B). For brevity, we refer to individual electrode contacts as *electrodes* through the rest of this paper. The movie clips presented to patients here were also used in prior work by our lab to uncover selectivity for acoustic and phonological information in the brain using scalp-EEG (Desai et al., 2023, 2024, 2021).

**Figure 1:**
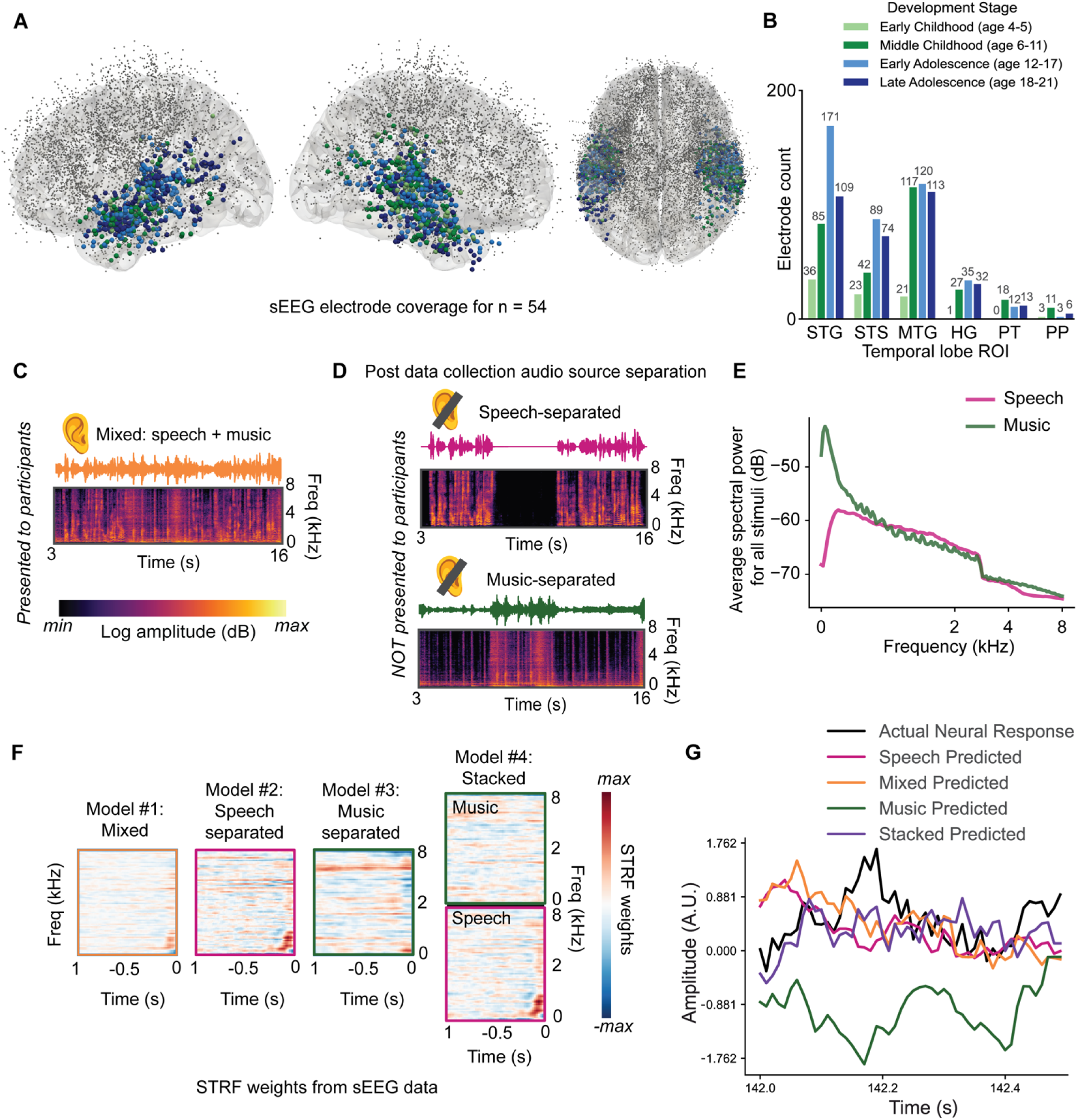
Framework for modeling intracranial responses to mixed and DNN-separated speech and music stimuli. A. Electrode coverage across 54 sEEG patients projected onto the cvs_avg35_inMNI152 atlas surface. Electrode colors in the temporal lobe represent development stages (early childhood to late adolescence). Coverage was unilateral or bilateral depending on the patient. Gray electrodes were recorded outside of temporal lobe regions of interest and not included in analyses. B. Electrode count by age (bars) and temporal lobe ROIs: superior temporal gyrus (STG), superior temporal sulcus (STS), middle temporal gyrus (MTG), Heschl’s gyrus (HG), planum temporale (PT), planum polare (PP). Bar colors indicate development stages: early childhood (ages 4-5, light green), middle childhood (ages 6-11, dark green), early adolescence (ages 12-17, light blue), late adolescence (ages 18-21, dark blue). C. Original mixed stimulus containing overlapping speech and music, shown with waveform and spectrogram. This represents the actual audio played to participants. D. Post hoc separation of the mixed (orange) audio into its isolated speech-separated (pink waveform) and music-separated (green waveform) components for computational modeling. These were not heard by the participant. E. Average spectral power (dB) of the speech and music signals for all stimuli in the study, computed from Mel spectrograms (80 Mel bands, 0-8 kHz). F. Spectrotemporal receptive field (STRF) encoding models were trained separately for each condition: mixed (orange), speech-separated (pink), music-separated (green) and stacked. Example STRF weight matrices are shown for one electrode. These weights represent which combination of acoustic features occurring in the past are most likely to increase the high gamma response (70–150 Hz) at time t=0. G. Example simulated model predictions of the neural activity for each condition, compared to the original (actual) brain response (black) from when sEEG patients listened to the mixed stimulus. In this example, the actual held out brain data is best modeled by the speech-separated spectrogram features (pink), and not well-modeled by the music-separated features (green).

Using deep neural network-based source separation (see Methods), we decomposed the mixed audio (Figure 1C) into constituent speech and music components (Figure 1D) and fit spectrotemporal receptive field (STRF) encoding models to predict neural responses to the spectrogram feature. Critically, patients only watched and listened to the mixed (original) audio (Figure 1C) and not the separated versions. We compared four linear encoding models to characterize how spectrogram features from speech and music streams were represented in neural activity: (1) the mixed model, which uses spectrotemporal features from the original, complete stimulus that was heard and seen by the participant, (2) the speech-separated model, which uses neural network derived vocal information only, (3) the music-separated model, which contains instrumental music only, (4) the stacked model, which includes both speech and music features as separate predictors within a single framework (Figure 1F). (See also Supplemental audio files 1-3).

Each of these models represents a different hypothesis about how the brain encodes auditory information in a mixed stimulus. For example, does the brain faithfully encode frequency features across both streams of the mixed auditory stimulus, or do some brain areas respond to certain frequencies within speech or music only? If the speech-separated or music-separated models outperform the mixed (true audio) model, this could represent a neural mechanism for auditory streaming in natural environments. If the stacked model outperforms single stream models, this would suggest that distinct neural populations encode both streams simultaneously, a form of multiplexed auditory processing (Agmon et al., 2022; Kaufman & Zion Golumbic, 2023; Vanthornhout et al., 2019). For all models, we predicted the same high-gamma neural activity (70-150 Hz) recorded during mixed listening. Models differed only in their input features, allowing us to test which acoustic representation best explained observed brain activity. Model performance was operationalized as the linear correlation (reported as r-value) between held-out high gamma signals for new stimuli and the high gamma neural activity predicted by the model for each electrode (see Methods; Figure 1G for an example).

### A bias for speech in STG

We compared the mixed, speech-separated, music-separated, and stacked models across electrodes in the primary and nonprimary auditory cortex. Neural activity in the temporal lobe was generally explained more accurately by models incorporating speech information rather than music, revealing a widespread cortical preference for speech encoding (Figure 2). To compare the relative contributions of these models to predict performance, we calculated a selectivity index that represents the degree to which neural activity for a particular electrode was better predicted by the speech-separated model (pink) or the music-separated model (green), operationalized as (unique R^2^_speech_ – unique R^2^_music_)/(R^2^_stacked_), where unique R^2^_speech_ is (R^2^_stacked_ – R^2^_music_), and unique R^2^_music_ is (R^2^_stacked_ – R^2^_speech_). This index is bounded from –1 (more music selective) to 1 (more speech selective). The distribution of speech– and music-selective electrodes is shown on the brain plots in Figure 2, along with example receptive field weights for each of the four models in four electrodes, all within the superior temporal gyrus (STG). Electrodes e1 and e2 were most accurately predicted by the speech-separated and stacked models, while electrodes e3 and e4 showed weak and strong music selectivity, respectively.

**Figure 2:**
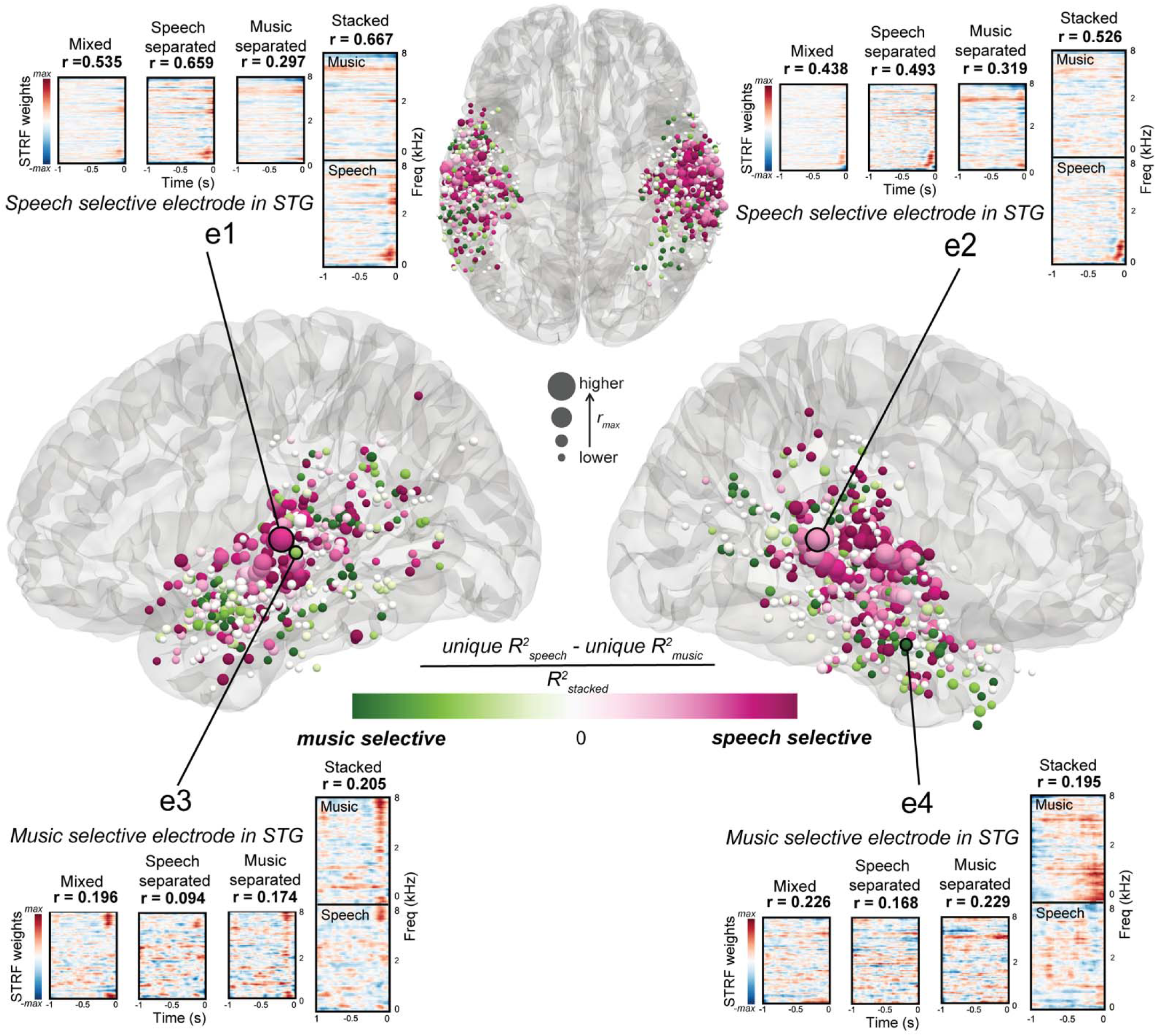
Superior temporal gyrus shows preference for speech over music in mixed stimuli. Electrode locations from 54 sEEG patients are projected onto the left and right hemisphere cortical surfaces. Electrodes are color-coded for selectivity – from most speech-selective (pink) to most music-selective (green), with electrodes with similar performance shown in white. Example STRF weight matrices are shown across time and frequency, with corresponding model performance (r) shown for representative speech selective (e1, e2) and music selective (e3, e4) electrodes. Larger electrode sizes represent better r values (actual vs. predicted maximum value for either model). STRF weights and model fits illustrate distinct encoding for speech vs. music, with stacked models capturing joint feature contributions. Electrode selectivity maps highlight functional organization of speech and music processing across the temporal lobe.

*Higher-order auditory cortex shows robust speech-selective encoding.* To quantify regional differences in model performance across the auditory cortex, we compared each model’s predictive accuracy within six temporal lobe regions of interest including primary (HG, PT) and nonprimary auditory cortex (STG, STS, MTG, PT) (Figure 3). In STG, speech-separated features significantly outperformed both music-separated and mixed features (Figure 3A). In STS and MTG, speech-separated features significantly outperformed music-separated features (Figure 3B-C). In all 3 higher-order regions (STG, STS, MTG), the stacked model, which jointly represents both speech and music features, significantly outperformed the mixed model and the music-separated model (Figure 3A-C and Supplemental Figure 1A-C). This suggests these regions have an overall bias toward speech-related acoustic features (Figure 2). Specific comparisons are detailed for each ROI and summarized in Table 1.

**Figure 3.**
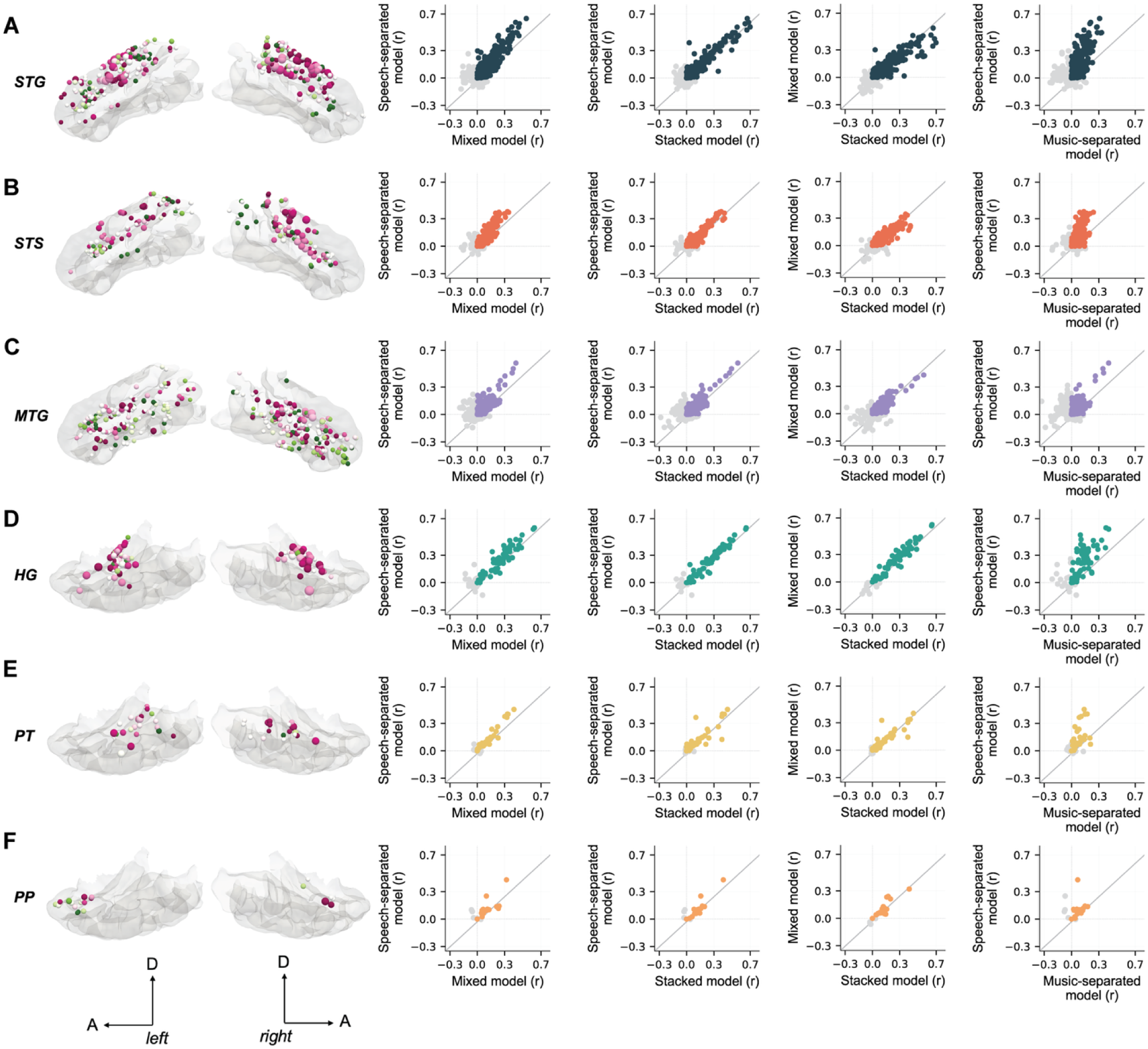
Higher-order temporal cortex shows selectivity for speech over mixed audio and music only. Brain plots show all participants’ electrodes projected onto a Montreal Neurological Institute (MNI) atlas brain (cvg_avg35_inMNI152). Electrodes are color-coded for selectivity – from most speech-selective (pink) to most music-selective (green) as in Figure 2. Larger electrode sizes on the brain represent better r values (larger maximum r). Scatter plots show speech-separated vs. mixed, speech-separated vs. stacked, mixed vs. stacked, and speech-separated vs. music-separated model comparisons. Each dot represents an electrode and dot color represents temporal lobe ROI. A. STG. B. STS. C. MTG. D. HG. E. PT. F. PP. Deviations from the unity line indicate which model better captures neural responses in each region, with points above the line indicating better performance for the y-axis model and points below the line indicating better performance for the x-axis model.

**Table 1:** Pairwise comparisons of model correlations between all models. Linear mixed-effects models comparing different models with model type, age, and their interaction as fixed effects, sex as a covariate, and random intercepts for subject. Negative β values indicate lower correlations for the second comparison condition relative to the first. STG = superior temporal gyrus; STS = superior temporal sulcus; MTG = middle temporal gyrus; HG = Heschl’s gyrus; PT = planum temporale; PP = planum polare. Significance levels: *p<0.05, **p<0.01, ***p<0.001.

| Region | Comparisons | $\beta$ | t(df) | p |
| --- | --- | --- | --- | --- |
| STG | **Speech-separated vs. Mixed** | **-0.022** | **-2.717(1198)** | **0.006\*\*** |
|  | Speech-separated vs. Stacked | 0.006 | 0.736(1196) | 0.461 |
|  | **Mixed vs. Stacked** | **0.029** | **3.453(1193)** | **0.0005\*\*\*** |
|  | **Speech-separated vs. Music-separated** | **-0.067** | **-7.760(1201)** | **$1.81 \times 10^{-14}$ \*\*\*** |
| STS | Speech-separated vs. Mixed | -0.015 | -1.945(685) | 0.052 |
|  | Speech-separated vs. Stacked | 0.011 | 1.499(684) | 0.134 |
|  | **Mixed vs. Stacked** | **0.027** | **3.427(683)** | **0.0006\*\*\*** |
|  | **Speech-separated vs. Music-separated** | **-0.039** | **-4.4880(686)** | **$1.32 \times 10^{-6}$ \*\*\*** |
| MTG | Speech-separated vs. Mixed | -0.008 | -1.345(1044) | 0.178 |
|  | Speech-separated vs. Stacked | 0.007 | 1.299(1041) | 0.194 |
|  | **Mixed vs. Stacked** | **0.015** | **2.612(1033)** | **0.009\*** |
|  | **Speech-separated vs. Music-separated** | **-0.022** | **-3.673(1051)** | **0.0002\*\*\*** |
| HG | Speech-separated vs. Mixed | 0.007 | 0.442(295) | 0.658 |
|  | Speech-separated vs. Stacked | 0.019 | 1.135(295) | 0.257 |
|  | Mixed vs. Stacked | 0.011 | 0.678(295) | 0.498 |
|  | **Speech-separated vs. Music-separated** | **-0.095** | **-5.419(295)** | **$1.24 \times 10^{-7}$ \*\*\*** |
| PT | Speech-separated vs. Mixed | -0.018 | -0.998(126) | 0.320 |
|  | Speech-separated vs. Stacked | 0.009 | 0.511(126) | 0.610 |
|  | Mixed vs. Stacked | 0.027 | 1.506(126) | 0.134 |
|  | **Speech-separated vs. Music-separated** | **-0.067** | **-3.539(126)** | **0.0005\*\*\*** |
| PP | Speech-separated vs. Mixed | 0.016 | 0.766(66) | 0.446 |
|  | Speech-separated vs. Stacked | 0.026 | 1.258(66) | 0.212 |
|  | Mixed vs. Stacked | 0.010 | 0.479(65) | 0.633 |
|  | Speech-separated vs. Music-separated | -0.030 | -1.446(66) | 0.152 |

*Lack of speech enhancement in primary auditory cortex.* In the primary auditory cortex in HG (Figure 3D), neural activity was significantly less predictable by music-separated features than by any other model. However, this area showed no preferential response to speech over the mixed or the stacked model (Table 1). This pattern was similar in nearby temporal plane regions PT and PP (Figure 3E-F), though interpretation is limited by a smaller electrode sample size (PT: n=43; PP: n=23; Table 1). This pattern of reduced performance for music-separated models without the corresponding improvement for speech-separated models suggests that HG and temporal plane areas do not exhibit the same degree of stream-specific tuning as downstream regions, consistent with their role in early auditory processing. This absence of speech enhancement in HG, combined with strong speech selectivity in STG, suggests that content-specific filtering emerges outside of primary auditory areas and may instead be performed by higher-order regions.

*Hemispheric considerations and electrode coverage.* Speech and music processing have been reported to show hemispheric asymmetries, with left hemisphere regions often showing stronger speech selectivity and right hemisphere regions showing music preferences (Peretz, 2006; Zatorre, Belin, et al., 2002). Because electrode placement in our cohort was determined by clinical needs, coverage varied widely across both participants, some had unilateral, while others had bilateral coverage. In addition, model performance differed within individuals. To better understand how speech and music selectivity may differ between hemispheres across development, future studies will need more balanced, bilateral electrode coverage.

Having established these regional patterns, we next examined how speech selectivity evolves across development from childhood through young adulthood.

### Speech selectivity strengthens with age in STG

We next examined whether and how the encoding biases observed across different auditory regions change across childhood and adolescence. We were interested in the degree to which this bias towards speech selectivity might strengthen across age.

To examine developmental effects, we fit linear mixed-effects models to investigate the influence of age, sex, and model type (mixed, speech-separated, music-separated, stacked) on encoding model performance in temporal lobe regions (see Methods). STG showed significant developmental increases in speech encoding (Figure 4A). In this region, speech-separated models significantly outperformed both mixed models (β=-0.022; t(1198)=-2.711; p=0.006) and music-separated models (β=0.067; t(1201)=7.760; p<0.0001), demonstrating a clear speech bias. Critically, this speech preference strengthened with age: we observed a significant interaction between model type (music-separated vs. speech-separated) and age (β=0.045; t(1198)=2.370; p=0.017), indicating that STG becomes increasingly selective for speech over music with development (Figure 4A). The stacked model, which included both speech and music predictors, performed comparably to the speech-separated model (β=0.006; t(1196)=0.736; p=0.461), confirming that even when both streams are available, STG preferentially represents the speech component. Consistent with this, the representative electrodes in Figure 2 with the strongest encoding are from early and late adolescence development groups (e1:15yo; e2:18yo; e3:21yo; e4:15yo); younger ages did not show robust representation of speech or music, consistent with the developmental trajectory observed in our statistical models.

**Figure 4.**
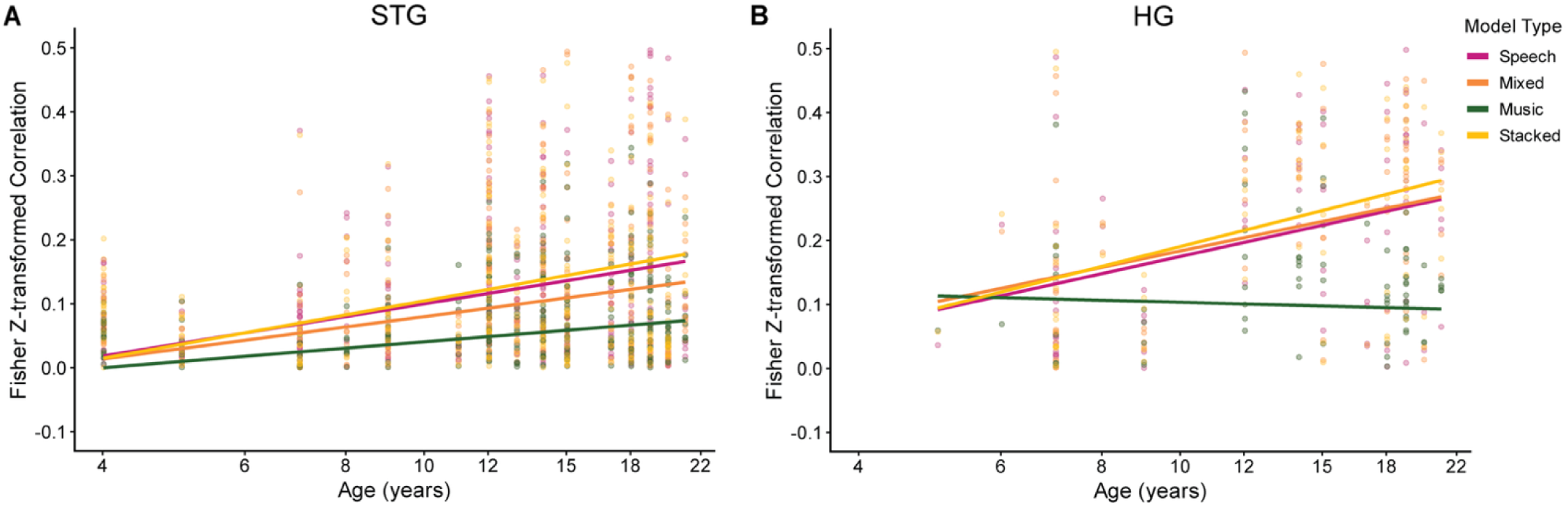
Developmental strengthening of speech selectivity in STG but not HG. Fisher Z-transformed encoding model correlations as a function of age (years, log-scaled and mean-centered) for four model types: speech-separated (pink), mixed (orange), music-separated (green), and stacked (yellow). Solid lines show linear regression fits for each group. Each dot represents one electrode. A. STG. Speech-separated models outperform both mixed and music-separated models, and this speech selectivity grows with age, reflecting progressive specialization for speech in higher-order auditory cortex. B. HG. Although a significant age x model type interaction is present, speech-separated models do not outperform mixed or stacked models at any age. The apparent developmental effect in HG instead reflects a relative decline in music-separated model performance with age, not preferential speech encoding, indicating that primary auditory cortex does not develop the same content-specific filtering observed in STG.

We next asked whether this developmental pattern was unique to STG by examining age interactions across the remaining temporal lobe regions. STS and MTG showed no significant age interactions (STS: all p>0.105; MTG: all p>0.786), indicating that while these regions may show speech selectivity, their encoding preferences do not change across the age range we sampled. In HG, however, we observed a significant age x model-type interaction (β=-0.153; t(295)=-3.617; p=0.0003; Figure 4B), but speech-separated models did not outperform mixed or stacked models at any age (speech-separated vs. mixed: β=0.007; t(295)=0.442; p=0.658; speech-separated vs. stacked: β=0.019; t(295)=1.135; p=0.257), indicating that HG does not show preferential speech encoding at any stage of development. In this case, a separate effect emerged—a relative decrease in performance of the music-separated models.

Together, these findings suggest that STG possesses specialized mechanisms for prioritizing speech information over music in complex auditory soundscapes, even without explicit attention to either stream, and that these selective processing mechanisms are established early but continue to refine throughout childhood and adolescence. In HG, while a significant age interaction was also observed, this does not reflect preferential speech encoding at any stage of development. No significant sex differences were observed in any regions.

*Effects of musical expertise.* Prior work has suggested that experience can modulate cortical entrainment to music (Doelling & Poeppel, 2015), but also that musical selectivity emerges without explicit training (Boebinger et al., 2021). To test these effects in our patient cohort, we investigated the relationship between musical training and speech vs. music selectivity. We collected survey data from a subset of N=19 participants on their musical background, including whether they play or have played a musical instrument, what instrument(s) they learned, duration of training, and frequency of musical activity. Among the respondents, 9 participants reported any musical training history while 10 reported no training. Among those with training, 6 participants reported regular musical practice (daily to several times per month), 2 participants reported having musical training but practicing irregularly or not at all, 1 did not report the practice frequency. Instruments included piano (N=2), voice/singing (N=2), percussion including drums and xylophone (N=2), guitar (N=2), and single reports of violin, recorder, harmonica, and keyboard. Training duration varied substantially, ranging from school-based instruction (reported as “at school” or specific grade levels) to extensive training reported as “since birth” or “since 2 years approx.” There was no significant difference in age between participants with musical training (mean=12.8 years, SD=4.7, range=7-19 years) and those without musical training (mean=12.4 years, SD=5.6, range=6-21 years; Mann-Whitney U=44.5, p=1.00).

To examine whether musical training influenced speech vs. music encoding, we conducted linear mixed-effects analyses on electrode-level model performance data from higher-order auditory regions (STG, STS, MTG) where speech selectivity was most prominent (see methods). Figure 5 shows Fisher Z-transformed model performance for participants with (purple) and without (green) musical training. Visually, both trained and untrained groups showed similar distributions for speech representation (Figure 5A), with most electrodes showing positive z(r_speech) values that increased modestly with age regardless of training status. In contrast, music representation (Figure 5B) was lower overall and showed flatter age trajectories in both groups, with considerable overlap between trained and untrained participants across the age range sampled.

**Figure 5.**
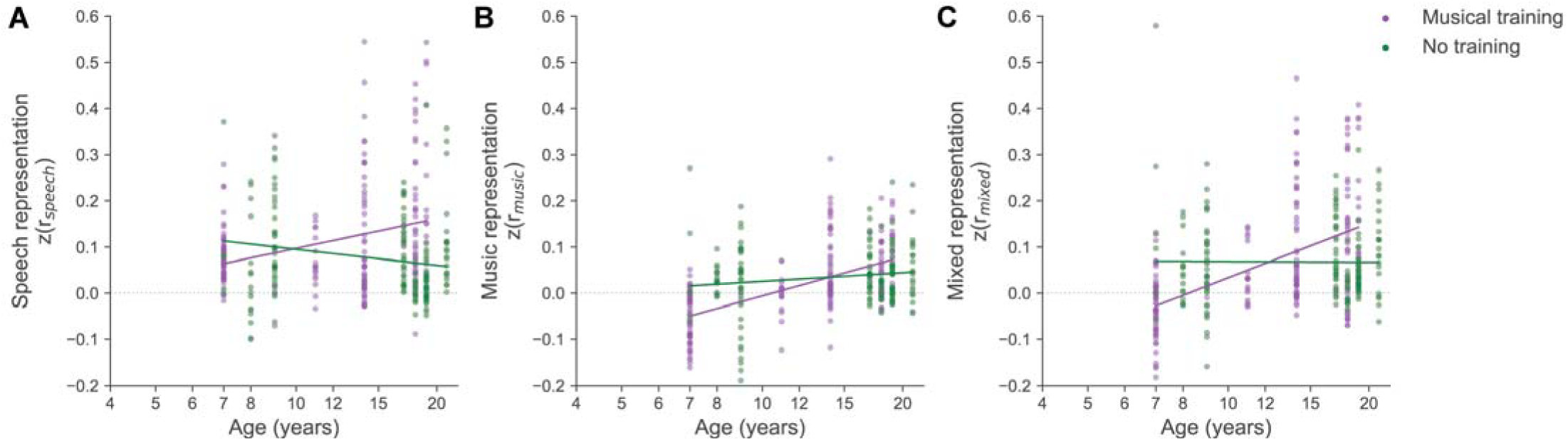
Speech and music encoding in higher-order auditory cortex as a function of musical training and age. Scatter plots show Fisher Z-transformed encoding model performance (r) for A. the speech-separated model, and B. the music-separated model, C. the mixed model across age (years, log-scaled), separately for participants with musical training (purple dots) and those without (green dots). Dashed lines show linear regression fits for each group. Data are from electrodes in higher-order auditory regions (STG, STS, MTG). Each dot represents one electrode.

Linear mixed-effects models revealed a significant two-way interaction between musical training and age (β=0.013; t(23)=3.498; p<0.001), indicating that musically trained participants have greater mixed model performance as they get older. We also observed a significant interaction between speech-separated model type and musical training (β=0.030; t(1443)=2.107; p=0.035), suggesting that trained participants’ neural responses in higher-order auditory cortex were better captured by isolated speech features relative to the full mixture, compared to untrained participants. Notably, no corresponding interaction was observed for music-separated models (β=-0.018; t(1443)=-1.260; p=0.208), suggesting that training did not enhance music-specific encoding, and that even in trained listeners, speech remained the dominant represented stream. A significant three-way interaction between music-separated model type, musical training, and age (β=-0.005; t(1443)=-1.970; p=0.049), suggesting that the separation between mixed and music model increases with age and musical training. No other significant three-way interactions emerged (all p>0.05). However, given the cross-sectional nature of our data and the relatively small sample size (N=9 and N=10 per group), we cannot draw strong conclusions about training effects on neural encoding independent of developmental maturation. Nevertheless, at a descriptive level, the similar distributions visible in both groups (Figure 5A-C) indicate that the fundamental speech bias observed in higher-order auditory cortex, wherein speech-separated models outperform music-separated models, emerges regardless of musical training status, consistent with prior findings that music selectivity does not require explicit training (Boebinger et al., 2021).

## Discussion

Despite extensive research on cortical selectivity for speech and music, how the human auditory cortex encodes multiple overlapping sound sources during naturalistic listening, and how this process develops across childhood, has remained unclear. To address this gap, we used high spatiotemporal resolution sEEG to investigate neural responses to concurrent speech and music during passive, real-world listening. First, we found that higher-order cortical activity was better predicted by speech-separated features of the spectrogram, even when listeners heard a mixture of simultaneous speech and music. Second, this speech bias was strongest in higher-order auditory regions such as superior temporal gyrus (STG), superior temporal sulcus (STS), and middle temporal gyrus (MTG), but not in primary auditory cortex (HG). Third, age-related increases in speech selectivity are specific to STG and emerge across participants regardless of musical training. Remarkably, these effects emerged even though participants listened only to mixed stimuli containing both speech and music, suggesting that the auditory system spontaneously emphasizes speech-related information without explicit attention or task demands. Together, these findings indicate that, under passive listening conditions, the developing auditory cortex preferentially encodes speech, reflecting a hierarchical and maturing specialization for behaviorally relevant acoustic information.

*Segregation of sound in higher-order auditory cortex.* Our findings align with a model of auditory cortical organization in which early auditory regions represent spectrotemporal information, while speech-specificity is seen in higher-order auditory cortex (Hickok & Poeppel, 2007; Kaas & Hackett, 2000; Mesgarani et al., 2014; Rauschecker & Tian, 2000). Typically, posteromedial HG corresponds to primary auditory cortex (A1), characterized by precise tonotopy and the encoding of low-level spectrotemporal features (Brewer & Barton, 2016; Da Costa et al., 2015; Hamilton et al., 2021; Howard et al., 1996). By contrast, the STG and adjacent regions of STS and MTG show parallel phonological and linguistic selectivity (Chang et al., 2010; de Heer et al., 2017; Hamilton et al., 2021; S. V. Norman-Haignere et al., 2022). The stronger and developmentally increasing speech selectivity observed in STG likely reflects specialization for acoustic-phonological transformations that are uniquely important for speech perception. While HG primarily encodes the physical energy of sound mixtures (frequency, intensity, timing), high-order regions such as STG, STS, and MTG integrate over longer timescales and are tuned to amplitude and spectral modulations that match the structure of speech (Hullett et al., 2016; Jain et al., 2020; Lerner et al., 2011). Although electrode coverage in PT and PP was limited in our study, available data suggest that these regions may serve as intermediate stages linking low-level acoustic analysis with higher-order categorical processing (Angulo-Perkins et al., 2014; Timothy D. Griffiths & Warren, 2002), including processing accent variation (Adank et al., 2012). Electrical stimulation of PT has also been shown to reduce the perception of background sounds when mixed with speech, suggesting an important role in auditory streaming (P. Patel et al., 2022). A particularly notable result was that the speech-separated model outperformed the mixed model in higher-order cortex, even though participants only heard the mixed audio (Figure 3). This finding implies that the hierarchy in auditory cortex helps segregate and prioritize speech information within complex auditory mixtures, even without explicit attentional focus. The minimal added benefit of including music features in stacked models within STG and STS further suggests that music-related features contribute little to explaining neural activity in these regions during natural, speech-dominant listening.

*Developmental strengthening of speech selectivity.* Our developmental analyses revealed that speech selectivity strengthened with age, particularly in STG. This pattern aligns with behavioral evidence that children’s ability to segregate speech from background noise improves through adolescence (Calcus, 2024; Leibold, 2019; Sanes & Woolley, 2011; Vander Ghinst et al., 2019). Structural and functional maturation of higher-order association cortex, including changes in cortical thickness, myelination, and long-range connectivity proceeds more slowly than primary areas (Corrigan et al., 2021; Gogtay et al., 2004; Huttenlocher, 1979; Huttenlocher & Dabholkar, 1997; Lebel & Deoni, 2018). The confinement of both speech encoding and age effects to STG, rather than HG or STS, suggests regional heterogeneity in developmental trajectories: STG serves as an interface between acoustic and linguistic representations (Bhaya-Grossman & Chang, 2022; Bhaya-Grossman et al., 2026; Giordano et al., 2023; Hamilton et al., 2021; Mesgarani et al., 2014), and its protracted specialization may underpin emerging competence in language and communication. Even in younger participants, however, we observed measurable speech selectivity, suggesting that these cortical biases build upon early developing preferences for communicatively relevant sounds (Grossmann et al., 2010).

*Parallel but asymmetric encoding of speech and music.* While our analyses revealed a broad speech bias across much of the temporal lobe, we also observed distributed music related selectivity across the temporal lobe (Figure 3). However, these responses were weaker and less consistent across participants. This asymmetry aligns with prior fMRI and intracranial studies demonstrating that speech and music engage overlapping but partly distinct neural populations (Zatorre, Belin, et al., 2002), speech responses predominating in mid and posterior STG and STS, and music responses clustering in anterior STG and PP (S. Norman-Haignere et al., 2015; S. V. Norman-Haignere et al., 2022). Unlike speech selectivity, music selectivity did not increase across development and appeared increasingly dominated by the mixed model in older adolescents, suggesting a developmental trajectory distinct from speech. Importantly, this pattern cannot be attributed to stimulus imbalance: across all stimuli, the audio contained 9.9% isolated speech, 41.1% isolated music, 30.8% mixed (speech+music), and the remainder silent segments. Despite the substantial proportion of musical content, cortical activity across the temporal lobe was more accurately predicted by speech features, indicating an intrinsic speech bias even under acoustically rich and ecologically valid listening conditions. Interestingly, similar speech dominance effects have been reported in selective attention EEG and MEG studies, where participants actively attend to a sound (Ding & Simon, 2012; Kerlin et al., 2010; O’Sullivan et al., 2015; Zion Golumbic et al., 2013). Our findings extend these observations by demonstrating that a speech bias emerges even under passive listening, in the absence of explicit attentional cues. This speech dominance likely reflects the greater ecological salience and developmental importance of speech, which is encountered continuously and supports fundamental communication and learning processes from infancy, whereas music engagement is typically more variable and experience-dependent (Albouy et al., 2020; Peretz, 2006; Trainor & Corrigall, 2010).

*Speech and music selectivity in bilateral auditory cortex.* Across our participants, we observed speech-selective and weak music-selective responses across both left and right auditory cortex, though speech selectivity was considerably stronger and more widespread than music selectivity in both hemispheres (Figure 2). Primary auditory cortex in HG showed no preferential enhancement for speech-separated over mixed models, indicating that early auditory areas faithfully represent the complete acoustic input without stream-specific biases (Binder et al., 2000; T. D. Griffiths et al., 1998). In contrast, higher-order regions, including STG, STS, and MTG, which also encode detailed spectrotemporal information, showed robust speech selectivity bilaterally, with neural responses better predicted by speech-separated features than by the complete mixed spectrogram that the participants actually heard (Figure 2). This pattern suggests that content-specific filtering for speech emerges in bilateral association cortex, consistent with hierarchical models wherein phonological and lexical information is processed in both left and right temporal cortex, with connected speech recruiting additional left-lateralized networks (Peelle, 2012; Poeppel, 2003; Scott et al., 2000). The weaker music-selective responses were found in both temporal lobes, but they were less consistent and more variable across participants. This likely reflects the speech-dominant nature of our stimuli, as well as the passive listening task, which may have biased attention toward speech rather than music. The finding that both speech and music selectivity emerged bilaterally during passive movie watching, without explicit instructions to attend to either stream, suggests that auditory cortex automatically differentiates these acoustic categories, though the relative contributions of bottom-up sensory processing versus implicit attentional biases toward speech content remain to be determined (Davis & Johnsrude, 2003; Obleser et al., 2007).

*Unmixing auditory streams in natural listening.* A critical methodological advance in this study is the use of source separation combined with encoding models to probe the neural correlates of auditory scene analysis without manipulating attention or stimulus presentation. Unlike previous studies that relied on isolated or task-based stimuli (Belin et al., 2000; Boebinger et al., 2021; McCarty et al., 2023; S. Norman-Haignere et al., 2015), our participants listened passively to naturalistic audiovisual movie clips containing overlapping speech and music conditions that more closely replicate real-world listening. The finding that the brain’s responses to mixed stimuli were better predicted by isolated speech features supports the notion of automatic auditory stream segregation, a process long hypothesized in perceptual studies (Bregman, 1994; Cherry, 1953; Treisman, 1969) but rarely demonstrated with invasive neural data in developmental populations. These results also extend prior intracranial studies showing distinct oscillatory cortical tracking for speech and music (Osorio & Assaneo, 2025), by demonstrating that such differentiation occurs spontaneously and strengthens with age. The evidence for robust speech selectivity in STG, STS, and MTG, even under passive conditions, suggests that the brain continuously organizes complex soundscapes into meaningful streams, potentially forming the foundation for effortless communication in noisy environments.

*Implications.* Developmental strengthening of speech selectivity in STG may reflect the progressive refinement of neural circuits supporting language comprehension and social communication. Atypical maturation of these pathways could contribute to the speech-in-noise perception challenges often observed in developmental language disorders, dyslexia, auditory processing disorder, and autism (Dole et al., 2014; Ruiz Callejo et al., 2023). By characterizing how the cortex automatically prioritizes speech amidst competing sounds, these findings provide a neural framework for understanding individual differences in language learning and listening abilities. Beyond basic insight, delineating this functional pathway offers translational value for optimizing auditory training programs, tailoring speech therapies and music training in childhood, and guiding neurotechnological development, such as hearing devices and speech decoding algorithms that harness biologically grounded principles of cortical organization (Défossez et al., 2023; Lo et al., 2020; Neves et al., 2022; Nguyen et al., 2025).

*Limitations.* Although sEEG gives us unmatched temporal precision and direct access to local neural populations, electrode coverage was dictated by clinical requirements rather than experimental optimization, yielding uneven samples across individuals, cortical subregions, and development stages. In addition, limited data from our youngest age group (ages 4-5) temper the strength of early developmental inferences. We were also only able to collect data on musical training on a subset of participants (N=20). Beyond these methodological factors, the present linear encoding models capture only a subset of the transformations implemented by the auditory cortex. Incorporating nonlinear frameworks, such as deep neural networks constrained by biology, could expose higher-order representational structure and cross correspondence between brain and behavior. Furthermore, while we gave our participants no explicit instructions about whether to attend to the speech or the music, it is possible that in typical movie clips speech is more salient. Thus, future studies will incorporate direct attention and SNR manipulations to determine to what extent the speech bias we observed here can be reversed or tempered by explicit attention to music.

Despite these limitations, these findings reveal that the auditory cortex segregates and prioritizes speech over music during natural listening, with this capacity strengthening throughout development. This provides a neural mechanism for the ‘cocktail party’ phenomenon and establishes a foundation for understanding why children struggle more than adults to process speech in noisy environments.

## Methods

We conducted stereo-electroencephalography (sEEG) recordings on 54 individuals aged 4-21 years (29M/25F; ages 4-21; mean 12.59 ± 4.7 years) undergoing epilepsy monitoring at Dell Children’s Medical Center (DCMC) in Austin, TX and Texas Children’s Hospital (TCH) in Houston, TX. As part of their clinical evaluation, patients were implanted with depth electrodes (AdTech Spencer Probe Depth electrodes, 5mm spacing, 0.86mm diameter, 4-16 contacts per device), strip electrodes (AdTech), or grid electrodes (AdTech custom order, 5mm spacing, 8×8 contacts per device). At DCMC, PMT depth electrodes (5 mm spacing, 0.8mm diameter, 4-16 contacts per device) were also used. These electrodes covered auditory and language-related areas in either the right hemisphere, left hemisphere, or bilaterally. The clinical team determined the specific placement of electrodes based on the suspected origin of each patient’s seizures, but generally included broad coverage across the temporal, frontal, parietal, occipital lobes, insula, and limbic structures (Table 2). Full brain coverage is shown for all patients in Figure 1A. Electrodes with seizure-related activity were excluded from our analysis to avoid confounding factors related to pathological neural activity. We recorded neural activity during passive listening tasks in superior temporal gyrus (STG), middle temporal gyrus (MTG), superior temporal sulcus (STS), Heschl’s gyrus (HG), planum polare (PP), planum temporale (PT), and nearby temporal lobe regions. The University of Texas at Austin Institutional Review Board approved all procedures and research at both sites was performed under a single IRB agreement with UT Austin. Informed written consent and/or assent and parental permission were obtained from all patients or their families (for minor patients) prior to participation.

**Table 2:** Demographic and clinical characteristics of pediatric and adolescent patients undergoing stereo-electroencephalography (sEEG) for epilepsy monitoring. Data includes subject ID, age, sex, development stage, seizure lateralization and focus, and number of temporal lobe electrodes for all 54 patients (29M/25F; ages 4-21; mean 12.59 ± 4.7 years).

| Subject | Age | Sex | Development | Seizure Focus and Seizure Side | Number of Temporal Lobe Electrodes |
| --- | --- | --- | --- | --- | --- |
| S0005 | 15 | M | early adolescence | R parietal | 4 |
| S0006 | 14 | M | early adolescence | R hemisphere | 24 |
| S0009 | 12 | M | early adolescence | L frontal-temporal-parietal | 17 |
| S0011 | 19 | M | late adolescence | Bifrontal, R frontal-parietal | 38 |
| S0012 | 9 | M | middle childhood | Bilateral frontal | 26 |
| S0013 | 4 | F | early childhood | Bilateral midline | 28 |
| S0016 | 11 | M | middle childhood | R temporal | 13 |
| S0018 | 14 | F | early adolescence | L temporal | 20 |
| S0021 | 9 | F | middle childhood | L hemisphere | 3 |
| S0022 | 18 | M | late adolescence | R parietal-temporal | 17 |
| S0024 | 17 | F | early adolescence | L frontal | 1 |
| S0025 | 7 | M | middle childhood | L frontal-temporal | 12 |
| S0027 | 11 | M | middle childhood | L frontal-central | 9 |
| S0030 | 7 | M | middle childhood | L temporal | 12 |
| S0031 | 8 | F | middle childhood | R parietal-occipital | 18 |
| S0033 | 15 | M | early adolescence | R frontal-temporal-parietal | 9 |
| S0036 | 12 | M | early adolescence | L parieto-occipital | 12 |
| S0038 | 7 | F | middle childhood | R hemisphere | 38 |
| S0039 | 20 | M | late adolescence | L frontal-central | 15 |
| S0040 | 15 | F | early adolescence | L temporal-parietal | 22 |
| TCH2 | 7 | F | middle childhood | L frontal, bilateral temporal | 6 |
| TCH3 | 18 | F | late adolescence | L temporal | 28 |
| TCH5 | 8 | M | middle childhood | L temporal | 2 |
| TCH06 | 14 | F | early adolescence | R frontal-parietal | 25 |
| TCH7 | 5 | F | early childhood | L frontal | 7 |
| TCH11 | 11 | F | middle childhood | R parietal-occipital | 5 |
| TCH13 | 18 | F | late adolescence | L occipital | 20 |
| TCH14 | 19 | F | late adolescence | L temporal | 36 |
| TCH15 | 12 | M | early adolescence | R occipital-temporal | 18 |
| TCH16 | 13 | M | middle childhood | R parietal | 2 |
| TCH18 | 7 | M | middle childhood | L frontal | 18 |
| TCH19 | 15 | M | early adolescence | L frontal-central | 6 |
| TCH20 | 12 | M | middle childhood | Bilateral frontal | 13 |
| TCH22 | 9 | F | middle childhood | Bilateral occipital | 11 |
| TCH28 | 19 | F | late adolescence | Bilateral | 12 |
| TCH29 | 18 | F | late adolescence | L temporal-parietal | 37 |
| TCH30 | 21 | F | late adolescence | L temporal | 18 |
| TCH37 | 17 | M | early adolescence | L temporal-frontal | 24 |
| TCH38 | 17 | M | early adolescence | L frontal | 7 |
| TCH39 | 9 | M | middle childhood | Bilateral | 14 |
| TCH42 | 9 | F | middle childhood | R frontal | 7 |
| TCH49 | 9 | M | middle childhood | R temporal-parietal | 9 |
| TCH50 | 17 | F | early adolescence | L temporal | 17 |
| TCH51 | 6 | M | middle childhood | L frontal | 1 |
| TCH52 | 11 | F | middle childhood | L temporal-insular | 18 |
| TCH53 | 15 | M | early adolescence | L temporal-frontal | 19 |
| TCH56 | 5 | M | early childhood | Bilateral | 29 |
| TCH58 | 20 | F | late adolescence | L frontal | 12 |
| TCH61 | 14 | M | early adolescence | L frontal | 16 |
| TCH62 | 13 | M | middle childhood | R frontal | 16 |
| TCH64 | 12 | M | early adolescence | R frontal-parietal | 36 |
| TCH65 | 19 | F | late adolescence | L temporal | 35 |
| TCH66 | 4 | F | early childhood | Bilateral | 5 |
| TCH69 | 13 | F | middle childhood | Bilateral frontal | 11 |

### Electrode localization

Electrode localization was performed by co-registering each participant’s T1-weighted MRI scan with their CT scan using freesurfer and the Python package img_pipe (Hamilton et al., 2017). Three-dimensional reconstructions of the pial surface were generated using the participant’s T1 MRI in Freesurfer, and anatomical regions of interest (ROIs) for each electrode were identified using the Destrieux parcellation atlas (Dale et al., 1999; Destrieux et al., 2010). For cross-subject visualization, electrodes were non-linearly warped to the cvs_avg35_inMNI152 template (Dale et al., 1999) following the procedures detailed by (Hamilton et al., 2017). This nonlinear warping preserved the anatomical region of interest (ROI) of each electrode but did not maintain the exact geometry of individual devices, such as depth electrodes or grids. Anatomical ROIs were always determined from electrodes in the participant’s original native space.

### Experimental design and stimuli

Participants watched and listened to up to 25 movie trailers. Data were only included in the analysis if they listened to at least 6 movie trailers (approximately 12 minutes of data) (Desai et al., 2023). Across all trailers, the audio consisted of 9.9% isolated speech, 41.1% isolated music, and 30.8% mixed speech and music timepoints. This varied audio composition allowed us to examine neural responses to different auditory features. The training and testing data were split by movie trailer stimulus, with the test set comprising the same two movie trailers (Paddington and Inside Out).

#### Post hoc audio source separation

While all patients listened to only the original (mixed) audio, we performed audio source separation for use in encoding models (described below in *Encoding Model Framework*). Audio source separation was performed using the Moises software (https://moises.ai, https://github.com/moises-ai/). Moises employs deep learning architectures for music source separation, drawing on models like U-Net (Stoller et al., 2018) and Demucs (Défossez et al., 2019), which separate sources while preserving transient information and rhythmic structure in different streams such as vocals, drums, and bass. The Moises models are trained on both public dataset MUSCB18 (https://sigsep.github.io/datasets/musdb.html) and MoisesDB (https://github.com/moises-ai/moises-db), a proprietary dataset containing 240 multitrack songs across 12 genres, specifically designed to enable fine-grained separation beyond the traditional four stems separation (Pereira et al., 2023). To validate separation quality, we cross-checked results using MVSep (https://mvsep.com/en), which integrates the MDX-Net (Music Demixing Network) architecture for music demixing. Specifically, the MVSep-MDX23 model combines the Demucs4 and MDX-Net architectures (Solovyev et al., 2023). High-quality source separation is critical for this analysis, as poor separation would introduce cross-contamination between speech and music features, obscuring the brain’s true selectivity for each stream. We therefore employed state-of-the-art separation tools and validated results across multiple architectures. U-Net uses an encoder-decoder architecture to learn spectrogram masks (Stoller et al., 2018), Demucs operates directly in the waveform time domain to better capture transient and rhythmic information (Défossez et al., 2019), and MDX-Net employs a two-stream architecture that processes audio in both time and frequency domains simultaneously (Solovyev et al., 2023). By combining these complementary approaches, we minimized artifacts and ensured that our separated streams accurately represented isolated speech and music content.

#### Validation of speech separation

To validate the speech and music separation algorithms, we compared Moises to MVSEP by calculating the linear correlation between the speech-separated files computed by each algorithm for each movie trailer stimulus. The correlation values showed a high degree of similarity between the two sets of speech-separated files, confirming the consistency of the separation process across different algorithms. We also played each of the separated speech and music tracks and multiple authors (RKA, LSH) confirmed that the subjective quality of the track separation. Due to their high degree of similarity, Moises-separated tracks were used in subsequent analysis.

### Neural data preprocessing

For recordings at Dell Children’s Medical Center, we used a 256-channel PZ2 or PZ5 amplifier connected to an RZ2 digital acquisition system (Tucker-Davis Technologies, Alachua, FL, USA). At Texas Children’s Hospital, we employed the clinical Natus Quantum system (Natus Medical Incorporated, San Carlos, CA, USA). All recordings were sampled at 2048 Hz. The data were subsequently downsampled to 512 Hz for analysis. Local field potentials (LFPs) were extracted from each electrode. To minimize line noise, a 60 Hz notch filter with harmonics at 120 Hz and 180 Hz was applied. We manually identified and excluded bad channels based on power spectra that exhibited low or flat signals or significant artifacts. Additionally, we manually detected and excluded epileptiform activity from further analysis. After artifact rejection, we performed offline re-referencing using a common average reference across all electrodes. The neural activity was filtered in the high-gamma range (70–150 Hz) using an 8-band Hilbert transform (Mesgarani 2014; Hamilton et al 2018; Hamilton et al 2021) to extract the analytic amplitude, which has been shown to correlate with multi-unit neural activity (Ray & Maunsell, 2011). The resulting high-gamma signal was down-sampled to 100 Hz, log-transformed, and then z-scored relative to the mean and standard deviation of neural activity across each recording session to normalize the data for subsequent analyses.

### Encoding model framework

We compared four linear encoding models to characterize how acoustic features from movie trailers were represented in neural activity (Aertsen & Johannesma, 1981; Holdgraf et al., 2017; Theunissen et al., 2000), as measured by high-gamma power (70-150Hz) of the local field potential. Each model represents different hypotheses for how auditory information is encoded in the brain: (1) the mixed model, which uses features from the original stimulus that was heard and seen by the participant (an 80 mel frequency band spectrogram from the original sound), (2) the speech-separated model, which uses the source separated vocal information only (80 mel frequency bands), removing any instrumental or background noise that is not speech, (3) the music-separated model, which is similarly derived to (2) but contains instrumental music only (80 mel frequency bands), and (4) the stacked model, which had both speech-separated and music-separated features as a separate but joint input (80 mel frequency bands for speech, and a separate 80 mel frequency bands for music, for a total of 160 mel-band features). The first model assumes that all spectrotemporal information (speech, music, background sounds, and effects) is represented similarly by the brain; the second, that spectrotemporal information in speech is preferentially processed while music is ignored; the third assumes that spectral information in music is preferentially represented; and the fourth, that the brain encodes both speech and music features in parallel. Based on prior work showing increased performance of phonological feature models representing speech as compared to the full auditory signal (Desai et al., 2021), we hypothesized that models incorporating the speech-separated spectrograms would perform better than the mixed or the music-separated representations in speech preferring regions, like STG.

Critically, all four models were fit to the same neural data: the high gamma responses recorded while participants listened to the original mixed movie trailers. The models differ only in their input features (mixed, speech-separated, music-separated, or stacked), allowing us to test which acoustic representation best predicts the observed neural responses.

The general form of these encoding models can be expressed as:

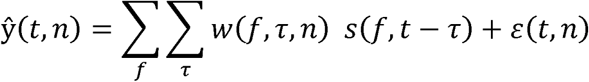

#### The mixed (speech+music) encoding model

The first model assumes that all acoustic information in the original movie trailer is encoded equally by the brain. Most importantly, this representation reflects the true stimulus that each participant heard. Here, 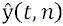 represents the predicted neural response for an electrode *n* at time *t*. *w*(f, τ, n) represents the learned spectrotemporal receptive field weights, estimated from neural data recorded when the participants listened to the mixed stimulus, for frequency bin *f* and time lag *r* at an electrode *n. s* represents the original stimulus spectrogram at frequency *f* and time lag r. ε(*t*, *n*) represents the residual error.

#### The speech-separated encoding model

In the second model, we were interested in whether neural activity recorded during mixed listening speech and music. Here, we use the same equation, but now *s*_speech_ represents the spectrogram is primarily driven by isolated speech components, even though the participant heard a mixture of of the isolated speech stream at frequency *f* and time lag *r*, obtained through the neural network-based source separation described in an above methods section.

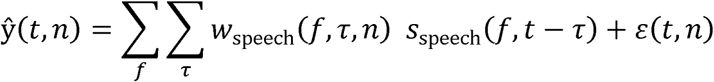

#### The music-separated encoding model

In the third model, we were interested in whether neural activity recorded during mixed listening was better predicted by isolated music components, even though the participant heard a mixture of speech and music. Here, *s*_music_ represents the spectrogram of the isolated music stream at frequency f and time lag r, obtained through neural network-based source separation described above.

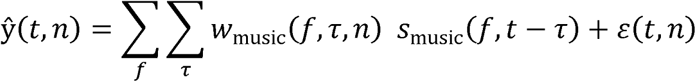

#### The stacked encoding model

In the fourth model, we tested whether neural responses recorded during mixed listening reflect multiplexed encoding of both speech and music features simultaneously. Rather than using a single spectrogram input, the stacked model combines both separated streams as independent predictors:

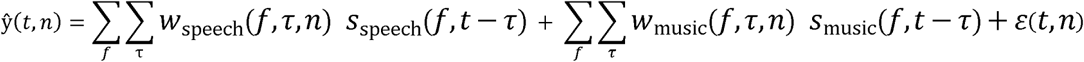

Here, *s*_speech_ and *s*_music_ represent separated speech and music spectrograms respectively, allowing us to assess how independently each sound stream contributes to neural activity when both are present simultaneously in the stimulus.

### Feature extraction and model fitting

For each model, we extracted spectrotemporal features using a mel-band approach with frequencies ranging from 0 Hz to 8 kHz, as in prior work (Hamilton et al., 2018; Mesgarani et al., 2014). We extracted 80 mel-frequency bands spaced logarithmically across this frequency range, using 25 ms windows with 10 ms step sizes, resulting in spectrograms sampled at 100 Hz. Both stimulus features and neural responses were z-scored across time prior to model fitting.

Spectrotemporal receptive fields were fitted considering time delays (lags) ranging from 0-1000 ms, a window sufficient to capture both speech-specific and music-related temporal dynamics. By incorporating multiple time lags as separate predictors, the model learns how acoustic features at different delays relative to the neural response contribute to brain activity, effectively characterizing the temporal structure of auditory processing.

Model weights were estimated using ridge regression with bootstrapped cross-validation to prevent overfitting and assess model stability. We tested ridge regularization parameters (alphas) from 10^2^ to 10^8^ in 20 logarithmic steps to determine optimal regularization for each electrode. For each bootstrap iteration (n=10), we randomly held out 3-second chunks of the training data (preserving temporal continuity to avoid autocorrection artifacts), fit the model on the remaining data for each alpha value, and evaluated performance on the held-out set using Pearson correlation between predicted and actual neural responses. The alpha value yielding the highest mean correlation across bootstraps was selected separately for each electrode. Final model weights were then estimated using the selected alpha on the full training dataset.

Model performance was quantified as a coefficient (r) between actual and predicted responses on an independent test set. This approach allowed us to examine how different acoustic features of naturalistic movie trailers are encoded in neural activity, with particular focus on whether the brain selectively represents speech vs. music information in complex auditory scenes.

### Noise ceiling estimation

To estimate an upper bound of our encoding model performance, we estimated noise ceilings from electrodes with repeated stimulus presentations (de Heer et al., 2017; Desai et al., 2021; Holdgraf et al., 2017; Pasley et al., 2012). The noise ceiling represents the maximum achievable correlation between predicted and actual neural responses given the inherent trial-to-trial variability in the neural signal. For each electrode with repeated presentations of the same movie trailer stimulus, we extracted the high gamma activity (70-150 Hz) time series for each repetition. We then computed pairwise Pearson correlations between all pairs of repetitions for each electrode (e.g., for 3 repetitions: correlation between repetition 1 and 2, 1 and 3, and 2 and 3) and averaged these pairwise correlations to obtain the noise ceiling estimate for that electrode-stimulus pair. However, we did not normalize our encoding model correlations by the noise ceiling for several reasons particularly relevant to developmental datasets. First, normalizing developmentally variable ceilings could conflate maturational effects in signal reliability with genuine changes in neural encoding, potentially obscuring true developmental effects in speech selectivity. Second, our repeated stimulus data were limited in scope; not all electrodes had sufficient repetitions. Third, and most importantly, our primary research questions focus on *relative* differences in model performance across conditions (speech-separated vs. music-separated vs. mixed vs. stacked) and across development, rather than absolute performance relative to a theoretical maximum. For these within, electrode comparisons, unnormalized correlations are appropriate because noise characteristics remain constant across model types. Regardless, across our electrodes with adequate repeated presentations, noise ceilings ranged up to r=0.58 in STG, STS, and MTG. Our best performing speech-separated models achieved correlations up to 0.60 in late adolescence, approaching these estimates.

### Statistical analysis

All statistical analyses were conducted in R Statistical Software (v4.1.2; R Core Team 2021) using linear mixed-effects (LME) models (lmerTest package – (Kuznetsova et al., 2017)). Before running statistical models, we excluded low-correlation data points where all four model types (mixed, speech-separated, music-separated and stacked) yielded correlations ≤ 0.0. Six anatomical regions were included in statistical analysis: HG, PT, PP, STG, STS, and MTG. Additionally, within each model type, electrodes with negative Fisher Z-transformed correlations were excluded prior to fitting linear mixed-effects models, to focus inference on electrodes showing meaningful neural encoding.

To better approximate normality, correlation values were Fisher Z-transformed using the hyperbolic arctangent function (atanh) prior to statistical analysis. Age was log-transformed and then centered (zero-meaned) to facilitate interpretation of interaction effects. Centering age ensures that main effects of model type reflect differences at the average age in our sample, rather than at age zero, which is particularly important when including interaction terms.

To assess model performance and developmental effects, we fit an LME model for each anatomical region. The model assessed the effects of model type (four levels: mixed, speech-separated, music-separated, stacked), centered log-transformed age (continuous), and sex as fixed effects, with a random intercept for subject ID to account for individual variability and repeated measures within subjects. The model included an interaction term between model type and age to test whether the effect of model type (i.e. the difference between encoding models) changed across development: In R: *lmer*(*Z*(*correlation*) ∼ *model type* × *age* + *sex* + (1|*subject ID*). This model tested whether different acoustic representations (mixed vs. separated streams vs. stacked) differentially predicted neural activity, and whether encoding strength changed with age.

To examine whether musical training influenced speech vs. music encoding, we conducted additional LME analyses on a subset of participants for whom musical training data were available. This analysis focused on electrodes in higher-order regions (STG, STS, MTG) where speech selectivity was most prominent. Musical training was coded as binary variable (trained vs. untrained) based on self-reported history of playing a musical instrument. The model included musical training status, model type, centered log transformed age, and their interactions as fixed effects, with subject specific random intercepts to account for the nested structure of multiple electrodes within participants. In R: *lmer*(*Z*(*correlation*) ∼ *model type* × *musical training* × *age* + *sex* + (1|*subject ID*). This model tested whether musical training modulated the relative encoding of speech vs. music features, and whether any training effects varied across development. Given the small sample size in this analysis, we interpret these data with caution.

## Data and code availability

Code for encoding model fitting, statistical analyses, and figure generation has been made available on a GitHub at https://github.com/HamiltonLabUT/Agravat2026_speechmusic. Patient data are not publicly available out of respect for patient privacy and consent but may be shared upon reasonable request to the corresponding author.

An interactive web-based visualization tool for exploring electrode locations and speech/music selectivity across participants of various ages is available at https://hamiltonlabut.github.io/SpeechMusicViewer/.

## Acknowledgements

This work was supported by the National Institute on Deafness and Other Communication Disorders (1R01-DC018579 to L.S.H.) and the Coleman Fung Foundation. The authors would like to thank Leah Du, G. Lynn Kurteff, Diego Mac-Auliffe, Alex Foox, and Elise Rickert for their help in data collection.

