## Supplemental Materials for "Human auditory cortex preferentially tracks speech over music without explicit attention"

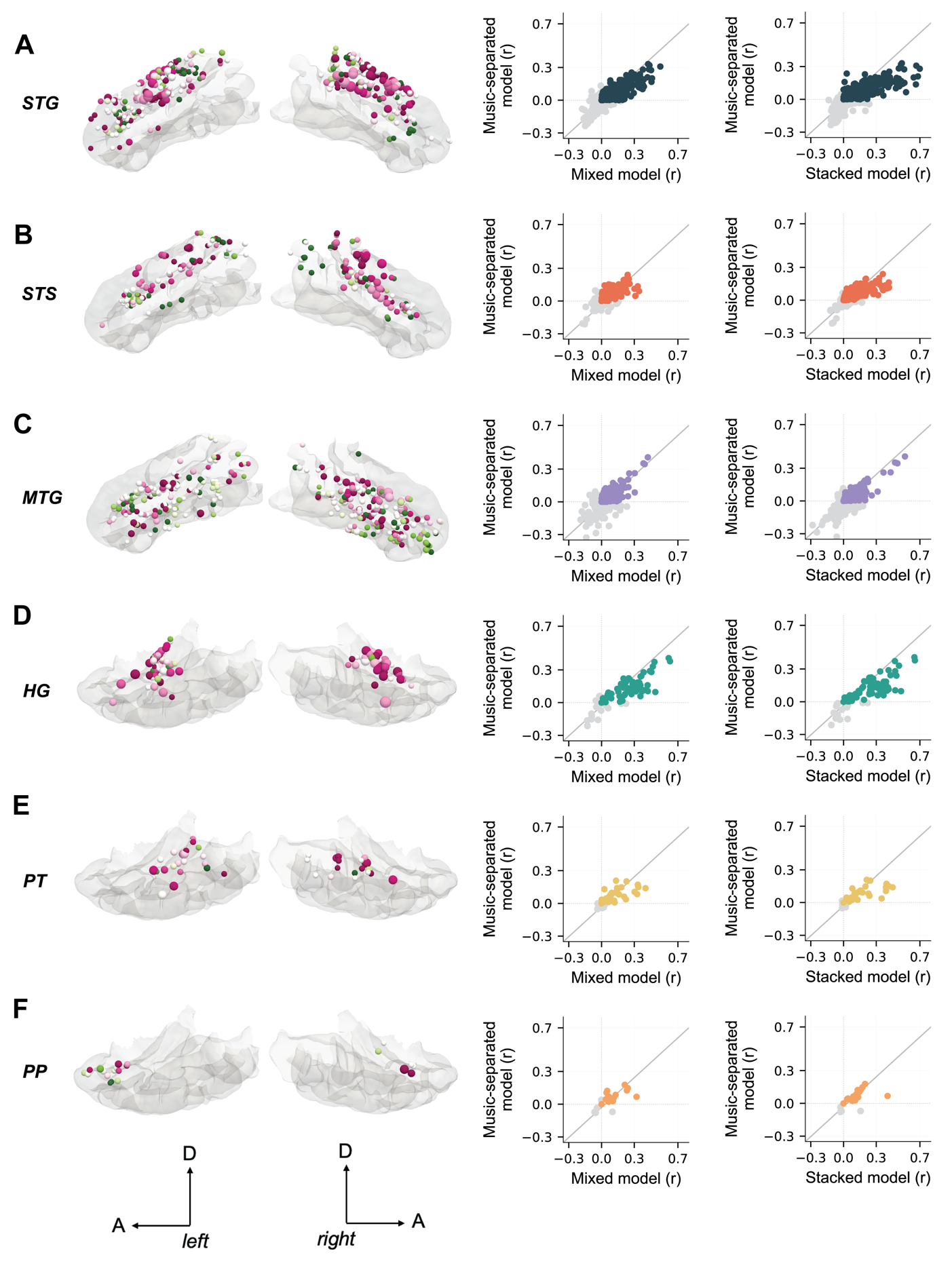
**Supplemental Information for *Agravat et al., 2026*.**

***Supplemental Figure 1.*** ***Model comparisons in higher order temporal cortex regions.*** *Scatter plots show music-separated vs. mixed, music-separated vs. stacked model comparisons. Each dot represents an electrode and dot color represents temporal lobe ROI. A. STG. B. STS. C. MTG. D. HG. E. PT. F. PP. Deviations from the unity line indicate which model approach better captures neural responses in each region, with points above the line indicating better performance for the y-axis model and points below the line indicating better performance for the x-axis model.*

***Supplemental Web Resource. Interactive Speech-Music visualization tool.*** *An interactive web-based tool for visualizing electrodes with varied speech and music selectivity among all the participants included in this study.* [*https://hamiltonlabut.github.io/SpeechMusicViewer/*](https://hamiltonlabut.github.io/SpeechMusicViewer/)
